# The evolutionary history of bears is shaped by gene flow across species

**DOI:** 10.1101/090126

**Authors:** Vikas Kumar, Fritjof Lammers, Tobias Bidon, Markus Pfenninger, Lydia Kolter, Maria A. Nilsson, Axel Janke

**Author notes:** Corresponding author: Axel Janke, Senckenberg Biodiversity and Climate Research Centre, Senckenberg Gesellschaft für, Naturforschung, Senckenberganlage 25, D-60325 Frankfurt am Main, Germany, email: Axel Janke.

## Abstract

Bears are iconic mammals with a complex evolutionary history. Natural bear hybrids and studies of few nuclear genes indicate that gene flow among bears may be more common than expected and not limited to the closely related polar and brown bears. Here we present a genome analysis of the bear family with representatives of all living species. Phylogenomic analyses of 869 mega base pairs divided into 18,621 genome fragments yielded a well-resolved coalescent species tree despite signals for extensive gene flow across species. However, genome analyses using three different statistical methods show that gene flow is not limited to closely related species pairs. Strong ancestral gene flow between the Asiatic black bear and the ancestor to polar, brown and American black bear explains numerous uncertainties in reconstructing the bear phylogeny. Gene flow across the bear clade may be mediated by intermediate species such as the geographically wide-spread brown bears leading to massive amounts of phylogenetic conflict. Genome-scale analyses lead to a more complete understanding of complex evolutionary processes. The increasing evidence for extensive inter-specific gene flow, found also in other animal species, necessitates shifting the attention from speciation processes achieving genome-wide reproductive isolation to the selective processes that maintain species divergence in the face of gene flow.

## Introduction

Ursine bears are the largest living terrestrial carnivores and have evolved during the last three million years, attaining a wide geographical distribution range (Fig. 1). Bears are a prominent case where conflicting gene trees and an ambiguous fossil record^1^ make the interpretation of their evolutionary history difficult^2^. Introgressive gene flow resulting from inter-species mating is believed to be rare among mammals^3^. However, some 600 mammalian hybrids are known^4^ and the importance of hybridization has started to gain attention in evolutionary biology^5^. Yet, our knowledge of the extent of post speciation gene flow is limited, because few genomes of closely related species have been sequenced.

**Figure 1:**
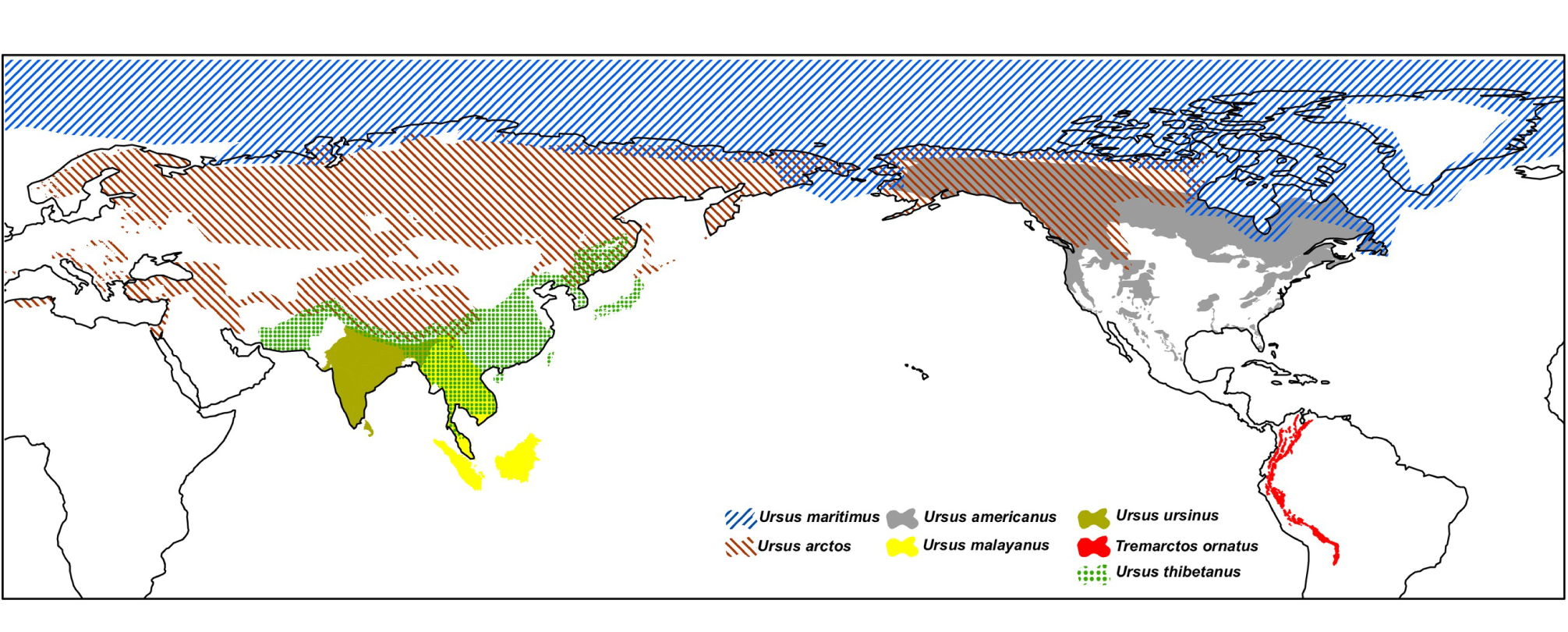
Approximate geographic distribution of extant bears according to IUCN data. Figure has been created using ArcGIS 10 (http://desktop.arcgis.com/en/arcmap/). The base map is obtained from the Global Administrative Areas (2012), GADM database of Global Administrative Areas, version 2.0 (www.gadm.org). Species range maps are from IUCN 2015 (www.iucnredlist.org).

In bears, natural mating between grizzlies (brown bears *Ursus arctos*), and polar bears (*Ursus maritimus*) results in hybrid offspring, the grolars^6,7^. Genome scale studies in brown and polar bears find that up to 8.8% of individual brown bear genomes have an origin in the polar bear^8^. Additionally, the brown bear mitochondrial (mt) genome was captured by polar bears during an ancient hybridization event^9,10^ and polar bear alleles are distributed across brown bear populations all over the world by male-biased migration and gene flow^8,11^.

Polar and brown bears belong to the sub-family Ursinae, which comprises six extant, morphological and ecological distinct species^12^, but hybridization among some ursine bears is possible. A natural hybrid has been reported also between the Asiatic black bear (*Ursus thibetanus*) and the sun bear (*Ursus malayanus*)^13^. In captivity more bear hybrids are known (between sun and sloth bear, Asiatic black and brown bear, and between American black and brown bear), some of them have been fertile^4^. Despite limited population sizes for most bears and apparently distinct habitats, morphology and ecology, molecular phylogenetic studies have been unable to unequivocally reconstruct the relationship among the six ursine bear species^2,14^. Especially, the evolution of the American (*Ursus americanus*) and Asiatic black bear is difficult to resolve, despite being geographically well separated (Fig. 1).

Evidence from the fossil record, morphology and mitochondrial phylogeny suggested a closer relationship between the Asiatic and the American black bears^14–17^. In contrast, autosomal and Y-chromosomal sequences support a different grouping with the American black bear being sister group to the brown/polar bear clade^2,11,14,18,19^. Another apparent conflict between mitogenomics, morphology and autosomal sequence data is the position of the morphologically distinct sloth bears (*Ursus ursinus*). Mitochondrial DNA (mtDNA) analyses and morphological studies placed sloth bears as sister-group to all other ursine bears, while nuclear gene analyses favor a position close to the sun bears^2,17,20^. A study of nuclear introns with multiple individuals for each ursine species was unable to reconstruct a well-supported species tree and suggested that incomplete lineage sorting (ILS) and/or gene flow caused the complexities in the ursine tree^2^. However, previous molecular studies did not have access to genome data from all bear species and were thus limited to small parts of the genome.

The genomic era allows for a detailed analyses of how gene flow from hybridization affects genomes, and has revealed much more complex evolutionary histories than previously anticipated for many species, including our own^21–24^. Multiple genomic studies on polar, brown bears and the giant panda^25–28^ lead to a wealth of available genomic data for some ursine bears. We investigated all living Ursinae and Tremarctinae bear species based on six newly sequenced bear genomes in addition to published genomes. Methods specifically developed to deal with complex genome data^29,30^ and gene flow in particular^21,31^ are applied to resolve and understand the processes that have shaped the evolution of bears.

## Results

The newly sequenced individuals were morphologically typical for the respective species. Mapping Illumina reads against the polar bear genome reference^28^ yielded for the Asiatic bear species an average coverage of 11X. Tables in (Supplementary Table 1 and 2) detail the sequencing and assembly data, and provide accession numbers of the included species. As a basis for subsequent evolutionary analyses, non-overlapping 100 kb **G**enome **F**ragments (GFs) were extracted from assemblies against polar bear scaffolds larger than 1 megabase (Mb). These have presumably a higher assembly quality than smaller fragments and still represent >96% of the genome (Supplementary Fig. 1). Ambiguous sites, gaps, repetitive sequences, and transposable element sequences were removed from GF alignments (Supplementary Fig. 2). The newly sequenced individuals were morphologically typical for the respective species. Pedigrees (Supplementary Fig. 3) and heterozygosity plots (Supplementary Fig. 4) show that the sequenced individuals are neither hybrids nor, compared to wild specimens, severely inbred.

## Network analysis depicts hidden conflict in the coalescent species tree

GFs larger than 25 kb, representing the majority of the length distribution (Supplementary Fig. 2), contain on average 104 substitutions between the Asiatic black bear and respectively, sun and sloth bear (Supplementary Fig. 5). Phylogenetic topology testing on real and simulated sequence data shows that GFs with this information content significantly reject alternative topologies (Supplementary Fig. 6 and Fig. 7). For subsequent coalescence, consensus, and network analyses only GFs larger than 25 kb were used and are thus based on firmly supported ML-analyses. A coalescent species tree utilizing 18,621 GFs >25 kb (869,313,834 bp) resolved the relationships among bears with significant support for all branches (Fig. 2A, Supplementary Fig. 8). In the coalescent-based species tree, sun and sloth bears form the sister group to the Asiatic black bear, and the American black bear groups with polar and brown bears. The spectacled bear (*Tremarctos ornatus*) is, consistent with previous results^2,18^, placed as sister taxon to Ursinae. The well-resolved coalescent species tree appears to be without conflict from the genomic data.

**Figure 2:**
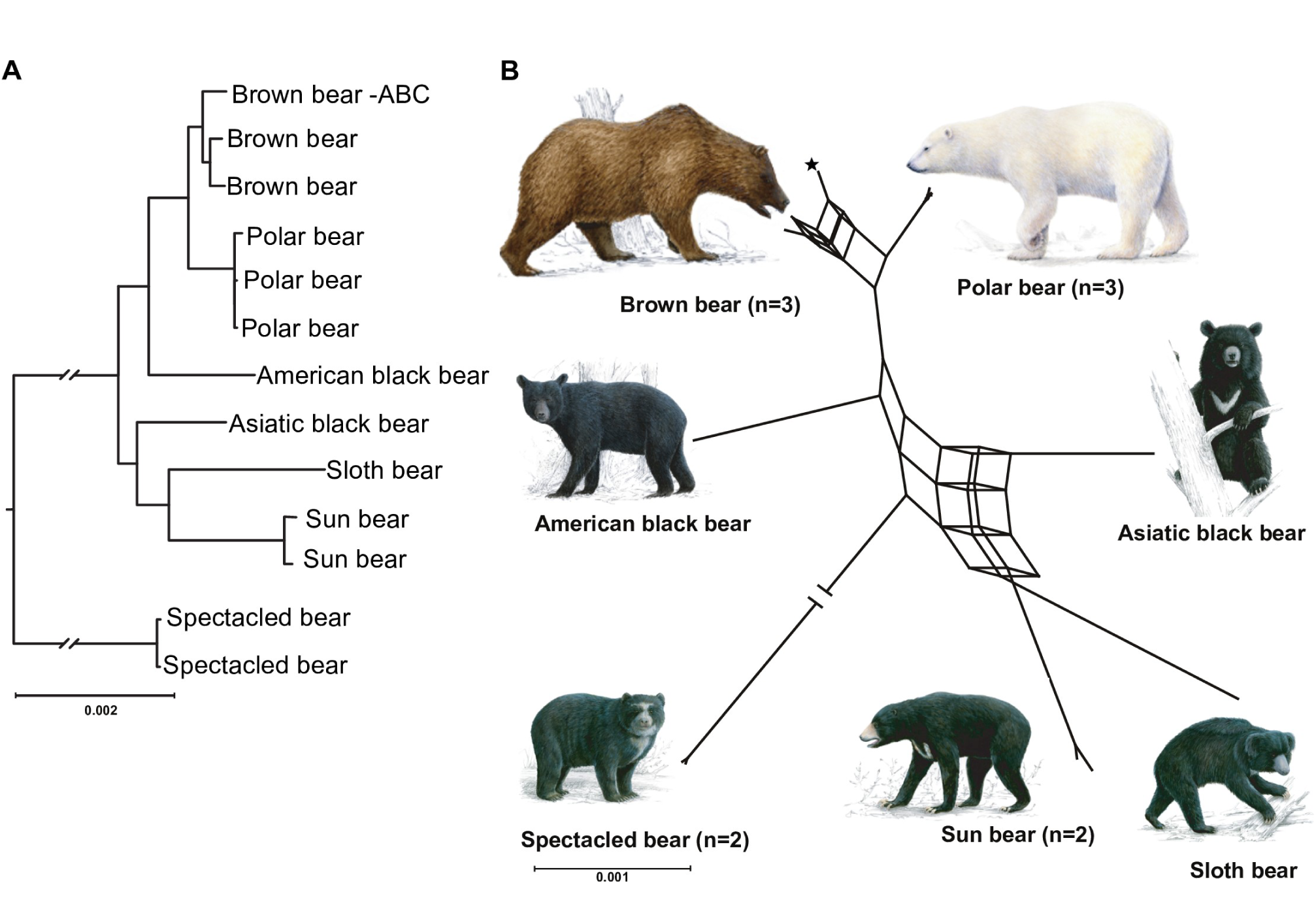
A coalescent species tree and a split network analysis from 18,621 GF ML trees. (A) In the coalescent species tree all branches receive 100% bootstrap support. The position of the root and the depicted branch lengths were calculated from 10 Mb of GF data. (B) A split network analysis with a 7% threshold level for conflicting branches depicts the complex phylogenetic signal found in the bear genome. The ABC-island brown bear (asterisk) shares alleles with polar bears; among the Asiatic bears allele sharing is more complex. The bear paintings were made by Jon Baldur Hlidberg (www.fauna.is).

However, a network analysis^32^ gained from the same 18,621 GFs identifies conflicting phylogenetic signal in the data (Fig. 2B). The square and cuboid-like structures are indicative of alternative phylogenetic signals, particularly among brown and polar bears, but also among the Asiatic bear species. The brown bear from the Admiralty, Baranof, and Chichagof (ABC) islands groups in different arrangements with the other brown and polar bears, consistent with gene flow between the two species^8,10,28^. When the threshold level for depicting conflicting branches is reduced in the network analysis, the signal becomes increasingly complex, illustrating the conflicting signals from the 18,621 ML-trees (Supplementary Fig. 9). Still, the network analysis agrees with the species tree when the spectacled bear is viewed as outgroup relative to the major cluster of American black, brown and polar bear, and Asiatic black bear with, sun and sloth bear. The phylogenetic conflict can be caused by incomplete lineage sorting (ILS) or gene flow, but less likely from lack of resolution due to the strong phylogenetic signal of each GF (Supplementary Fig. 6 and Fig. 7). Analyses of 8,050 protein coding sequences (10,303,323 bp) and GFs from scaffolds previously identified as X chromosomal (total 74Mb)^27^, also conform to the species tree and network (Supplementary Fig. 10). Finally, the paternal side of the bear evolution based on Y chromosome sequences^33^ cannot be analyzed for all species, but the resulting tree (Supplementary Fig. 11) is consistent with the inferred species tree.

The Bayesian mtDNA tree (Fig. 3, Supplementary Fig. 12) from 11,529 bp of sequence information that remained after alignment and filtering conforms to previous studies^2,17,34^, making this the hitherto largest taxonomic sampling of 38 complete bear mt genomes. However, several nodes of the mtDNA tree differ notably from the coalescent species tree (Fig. 2A). In the mtDNA tree, the brown bears are paraphyletic, because the brown bear mt genome introgressed into the polar bear population^10,35^. The extinct cave bear (*Ursus spelaeus*) is the sister group to polar and brown bears. The American black bear is the sister group to the Asiatic black bear, and the sloth bear is the sister group to all ursine bears. The topological agreement of the mtDNA tree to previous mtDNA studies and expected placement of the new individuals corroborates the matrilineal lineages and that the studied individuals are representative for their species and likely not hybrids.

**Figure 3:**
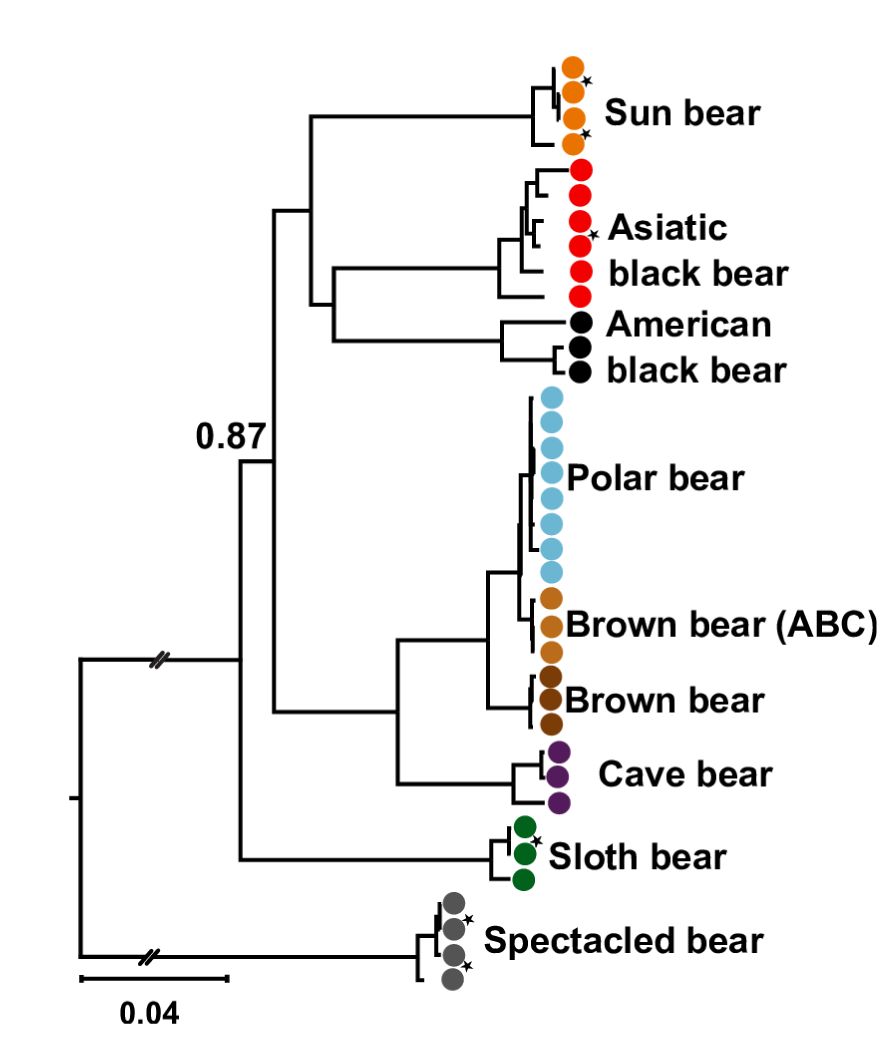
Phylogenetic relationship among the bears using mtDNA genomes. A Bayesian mitochondrial tree from 37 complete mt genomes (colored circles). Support values for p<1.0 are shown in (Supplementary Fig. 12) lists accession numbers. Stars indicate the mt genes that were sequenced in this study. Asterisks indicate new mt genomes and from the scale bar indicates 0.04 substitutions per site.

Finally, a consensus analysis based on ML-trees from GF (Supplementary Fig. 13) produces a tree that is identical to the coalescent species tree, but highlights that numerous individual GF trees support alternative topologies (Supplementary Table 3). Closer inspection of the individual 18,621 GF ML topologies shows that 38.1% (7,086) support a topology where Asiatic black bear is the sister group to the American black/brown/polar bear clade. The Asiatic black bear groups in different arrangements with the two other Asiatic bears: 18.7% (3,474) of the branches support a grouping with the sun bear, and 7.5% (1,394) with the sloth bear in other topologies.

## Gene flow among bears is common

Seemingly conflicting phylogenetic signals in evolutionary analyses can be explained either by incomplete lineage sorting (ILS)^36^ or gene flow among species. In contrast to the largely random process of ILS, gene flow produces a bias in the phylogenetic signal, because it is a directed process between certain species. The *D*-statistic measures the excess of shared polymorphisms of two closely related lineages with respect to a third lineage^21^ and can thus discriminate between gene flow and ILS. The test assumes that the ancestral population of the three in-group taxa was randomly mating and recently diverging^37^. These assumptions might be compromised in widespread, structured species like bears. However, speciation is rarely instantaneous, but is rather preceded by a period of population divergence. Such a period should not compromise the test as long as there was a panmictic population ancestral to the progenitor populations of the eventual daughter species at some point in time, which is a reasonable assumption.

The *D*-statistics analyses find evidence of gene flow between most sister bear species (Fig. 4, Supplementary Table 4-5 and Supplementary Fig. 14). Regardless if spectacled bear or giant panda is used as outgroup, the direction and relative signal strengths of gene flow in the tested topologies remain the same (Supplementary Table 6). The *D*-statistic is limited to four-taxon topologies and therefore gene flow signals are difficult to interpret when they occur between distant species, as it cannot determine if it is a direct, indirect, or ancestral signal. For taking more complex phylogenetic signals and gene flow patterns into account, and to determine the direction of gene flow, we applied the recently introduced *D_FOIL_*-statistics. This method uses a symmetric five-taxon topology and has specifically been developed to detect and differentiate gene flow signal among ancestral lineages^31^.

**Figure 4:**
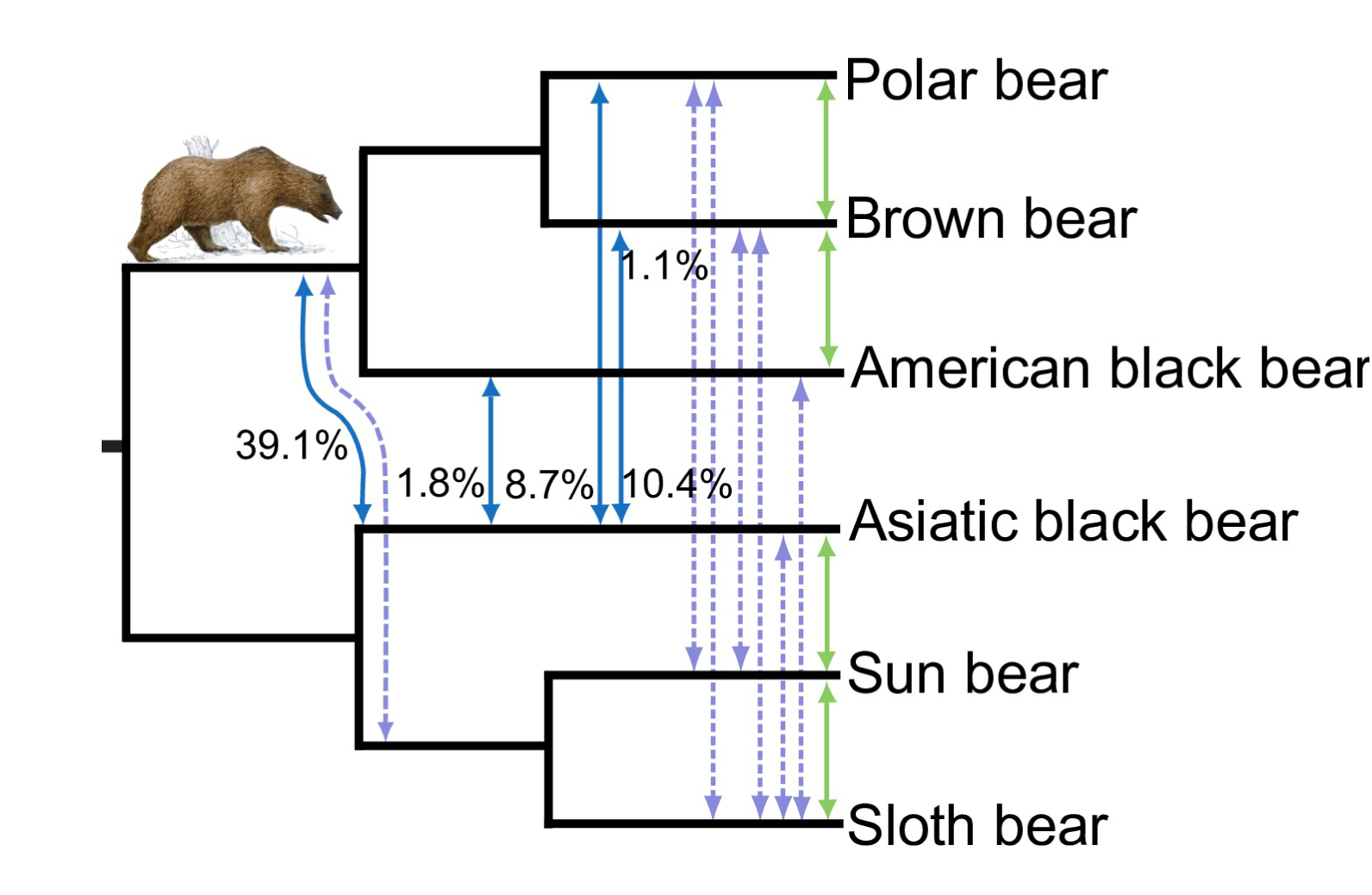
Graphical summary of gene flow analyses using *D* and *D_FOIL_* statistics on a cladogram. The percentage of GFs rejecting the species tree and indicating gene flow estimated by *D_FOIL_* analyses is shown by percentages and blue arrows for values >1%, and dashed lavender for <0.1% (Table 1). These percentages do not indicate the total amount of introgressed genetic material, which can be a fraction of the GF sequence. Green arrows depict *D*-statistics results for significant gene flow signal between recent species. Some gene flow cannot have occurred directly between species, because the involved species exist in different habitats. Therefore, it is suggested that such signals are likely remnants of ancestral gene flow or gene flow through a vector species. The bear painting was made by Jon Baldur Hlidberg (www.fauna.is).

In agreement with the phylogenetic conflict and *D*-statistics, the *D_FOIL_*- statistic finds strong gene flow between the ancestor of the American black bear/brown/polar bear clade, possibly the extinct Etruscan bear (*Ursus etruscus)*, and the Asiatic black bear (Fig. 4, Table 1). The Etruscan bear was geographically overlapping with other bear species and was, like the Asiatic black bear, widely distributed^38^. It has been identified in fossil layers of Europe between 2.5 Ma to 1.0 Ma^1,39,40^. The wide geographical distribution would explain the nearly equally strong gene flow from Asiatic black bear into brown bear also observed in the *D*-statistics analyses (Supplementary Fig. 14). Finally, there is a strong signal for gene flow between the American and Asiatic black bears. The gene flow could have taken place either on the American or Asiatic side of the Bering Strait and is consistent with the suggested mitochondrial capture between the species^2^ and the deviating mitochondrial tree (Fig. 3). It should be noted that most of the weaker gene flow signals in Fig. 4 (dashed-lines) need to be corroborated and do not necessarily reflect direct species hybridization. The signals are possibly remnants of ancestral gene flow that is not detected because of allelic loss or signal of indirect gene flow by ghost lineages or intermediate species. Permutations of species for the *D_FOIL_* analysis including other polar, sloth and brown bear individuals show that the estimates are taxon independent (Table 1).

PhyloNet^41^ has been specifically developed to detect hybridization events in genomic data while accounting for ILS. We applied the maximum likelihood approach implemented in PhyloNet^41,42^ to detect signs of hybridization among bear species. We sampled 4,000 ML trees from putatively independent GFs with one individual representing each species. The ABC island brown bear was chosen as another representative for brown bears and positive control, because for the ABC island brown bear population hybridization with polar bears has been documented^8,10,33^. The outgroup, the spectacled bears were removed to reduce the computational complexity and, because previous analyses could not detect gene flow between tremarctine and ursine bears. The complex phylogeny requires exceptional computational time so we analyzed only networks with up to two reticulations. The resulting PhyloNet network with the highest likelihood (Supplementary Fig. 15) shows reticulations between ABC island brown bear and polar bears, and also between the Asiatic black bear and the ancestral branch to American black, brown and polar bears. It is noteworthy, that the second reticulation has a high inheritance probability (41.8%), which agrees with the strongest gene flow signal identified by D_*FOIL*_ analyses (Fig. 4, Table 1). Due to computational limits so far only two reticulations that represent the strongest hybridization signals could be identified. For three and more reticulations the network-space becomes extremely large.

Additional analysis using CoalHMM^43^ supports the findings of gene flow from D-, *D_FOIL_*, and PhyloNet analyses (Supplementary Fig. 16). It shows that a migration model fits most pair wise comparisons significantly better than ILS. A sensitivity analysis shows that this finding is robust under a broad range of parameters (Supplementary Fig. 17 and 18). Thus, gene flow among bears throughout most of their history is the major factor for generating conflicting evolutionary signals.

## Estimation of divergence times and population splits

The phylogenomic based divergence time estimates (Fig. 5) are older than recent estimates based on nuclear gene data^2^, but are consistent with that on mtDNA data^17,44^ (Supplementary Table 7). The amount of heterozygous sites differs among species and individuals, and is highest in the Asiatic black bear genome and, as expected^2^ lowest in the polar bears and spectacled bears (Supplementary Fig. 4). It is noteworthy that the average numbers of heterozygous sites differ among the two sun bears, which may reflect different population histories.

**Figure 5:**
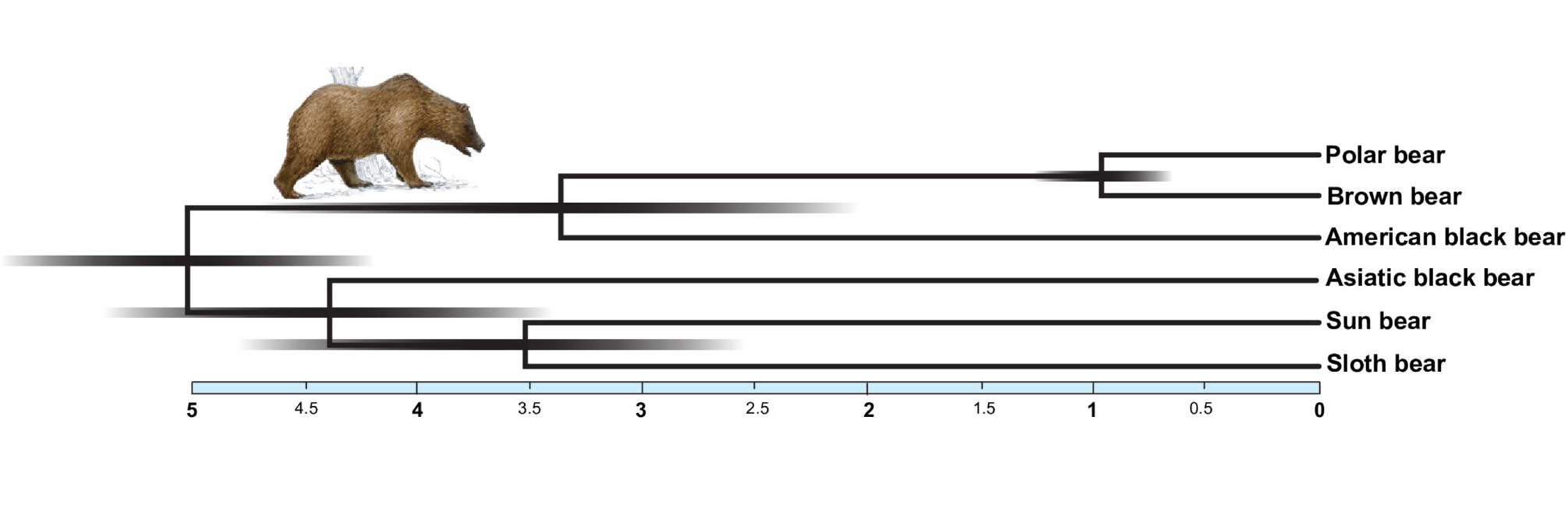
Phylogenomic estimates of divergence times. The light blue scale bar below the tree shows divergence times in million years and 95% confidence intervals for divergence times are shown at the nodes as shadings. Exact times and intervals for all divergences are listed in Supplementary Table 7. The bear painting was made by Jon Baldur Hlidberg (www.fauna.is).

Estimates for past changes in effective population size (*N*_*e*_) using the pairwise sequentially Markovian coalescent (PSMC)^45^ for all newly sequenced bears are shown in Fig. 6 (unscaled PSMC see Supplementary Fig. 19). While PSMC plots from low coverage genomes may vary and not be ultimately accurate, the plots inferred for the brown, polar and American black bear are very similar to previous published on higher coverage genomes (Supplementary Fig. 20)^26^. The demographic histories of the Asian bear individuals vary widely, but do not overlap in bootstrap analyses since 100 ka (Supplementary Fig. 21).

**Figure 6:**
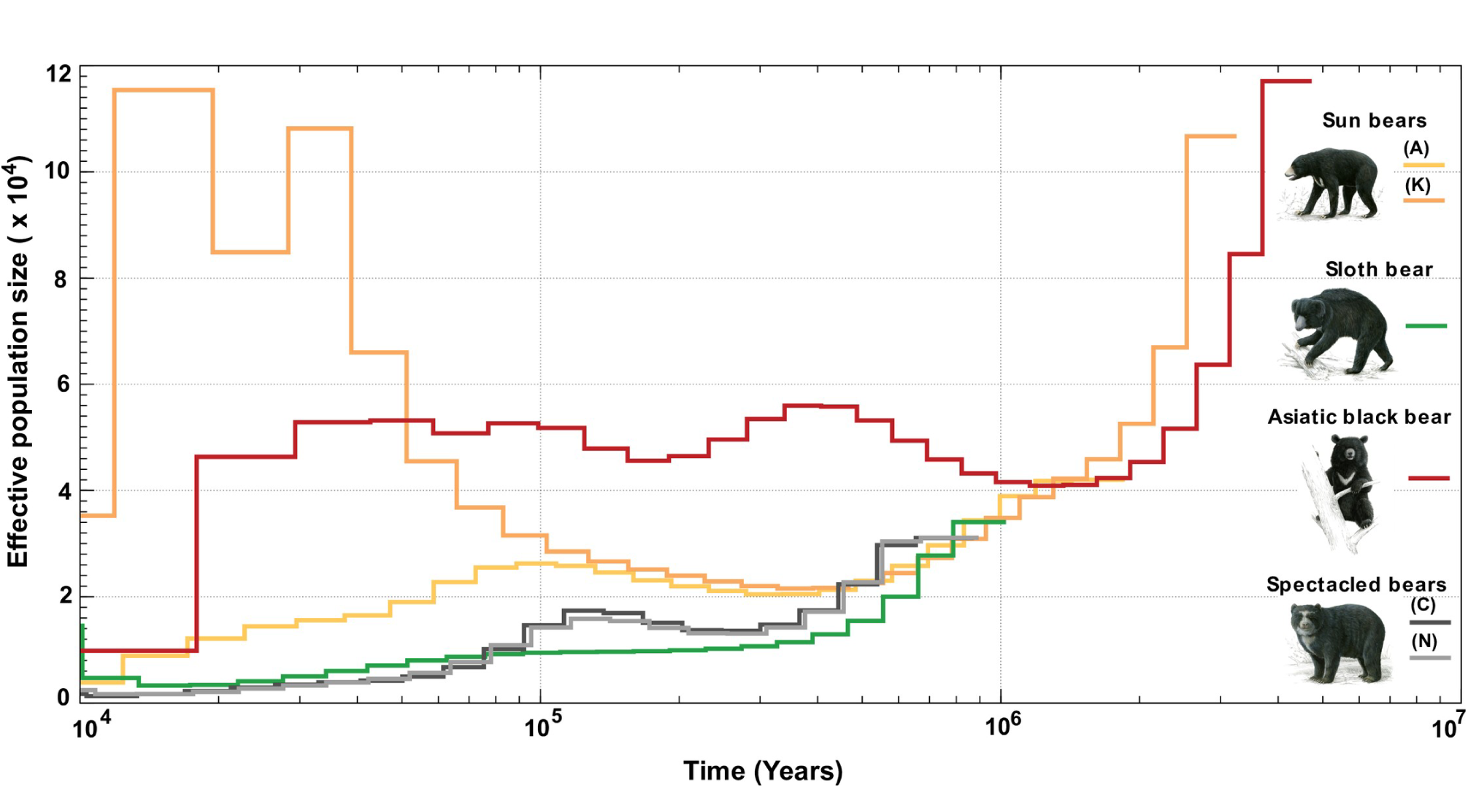
Historical effective population sizes (*N_e_*) using the pairwise Markovian coalescent (PSMC) analyses for the newly sequenced bear genomes. The x-axis shows the time and the yaxis shows the effective population size (*N_e_*). The two sun bears have radically different population histories that are not overlapping in bootstrap analyses since 100 ka (Supplementary Fig. 21) and the Asiatic black bear had a constant large Ne since 500 ka similar to that of the brown bear which is consistent with the wide geographic distribution and high heterozygosity. Other bears maintained a relatively low long-term Ne, consistent with a lower population diversity and low heterozygosity (Supplementary Fig. 4). The bear paintings were made by Jon Baldur Hlidberg (www.fauna.is).

According to the PSMC analyses the Asiatic black bear maintained a stable and a relatively high long-term *Ne* since 500 ka (Fig. 6). This is consistent with its wide geographic distribution and its high degree of heterozygous sites in the genome^2^. The effective population size of the Asiatic black bear declined some 20 ka, correlating with the end of the last ice age. By contrast, the spectacled bear maintained a relatively low long-term effective population size, consistent with their lower population diversity^2,46^. The demography of two sun bear individuals is strikingly different from each other since 100 ka. As the bootstrap replicates do not overlap, the different curves support a hypothesis of separate population dynamics (Supplementary Fig. 21). Their distinct mitochondrial lineages (Fig. 3) might indicate that the two sun bear individuals belong to the described subspecies *U. m. malayanus* (Sumatra and Asian mainland) and *U. m. euryspilus* (Borneo)^47^ respectively. The ancestor of extant sun bears might have settled in the Malay Archipelago during the marine isotope stage (MIS) 6. In the following Eemian interglacial, Borneo got isolated, thereby giving rise to different environmental conditions and to a distinct sun bear subspecies, but without samples from multiple individuals from known locations and high coverage genomes, this remains speculative.

## Discussion

Previously, nuclear gene trees and mitochondrial trees have been in conflict^17,18,34^, and a forest of gene trees made it difficult to conclusively reconstruct the relationships among bears^2^. Now, phylogenomic analyses resolve a solid coalescent species tree and provide a temporal frame of the evolutionary history of the charismatic ursine and tremarctine bears. This exemplifies the power of multi-species-coalescent methods that are becoming increasingly important in genomic analyses^48^. Phylogenetic networks show that evolutionary histories of numerous GFs, i.e. various regions of their genome, are significantly different. This is also evident from large-scale evolutionary analysis of insertion patterns of transposable elements into the bear genomes, which yield a similarly complex history bears^49^. However, phylogenetic networks show that evolutionary histories of numerous GFs, i.e. various regions of their genome, are significantly different. It is important to realize that bifurcating species trees, even coalescence based, can only convey a fraction of the evolutionary information contained in entire genomes and that network analyses are needed to identify underlying conflict in the data^29,50^. The analyses of the ursine phylogeny suggest that gene flow and not incomplete lineage sorting are the major cause for the reticulations in the evolutionary tree. These two processes can be distinguished from each other by methods and programs like *D-*statistics, *D_FOIL_* and Phylo-Net^21,31,41^ that are specifically developed for this task.

Some of the inferred gene flow between bear species appears weak or episodic and thus requires further corroboration by additional sampling of individuals. Population analyses show that American black bears are divided into two distinct clades that diverged long before the last glacial maximum, indicating a long and isolated evolutionary history on the North American continent^51^. Thus, it is unlikely that American black bears came into contact with the Asiatic sun and sloth bears^15,51^. Likewise, introgressive gene flow between south-east Asiatic bear species and polar bears requires an explanation, because they have been evolving in geographically and climatically distinct areas, from the time when polar bears diverged from brown bears and began parapatric speciation in the Arctic. It is therefore possible that some gene flow events occurred through an intermediate species. The brown bear has been shown to distribute polar bear alleles across its range^8^ and may therefore be a plausible vector species for the exchange of genetic material between Asiatic bears and the polar, or American black bear. The brown bear is a likely extant candidate, because it has been and is geographically wide-spread^52^. Furthermore, the geographical range of brown bears overlaps with all other ursine bear species (Fig. 1), they have reportedly migrated several times across continents and islands^52,53^, and numerous brown bear hybrids with other bears in either direction are known^4^. While also the Asiatic black bear was widely distributed across Asia and had, like the brown bear^26^, a large effective population size (Fig. 6), a migration of the Asiatic black bear into North America has not been shown. Likewise, migration of the American black bear in the opposite direction, from the American to the Asian continent, is not evident from fossil data. The D_*FOIL*_ and PhyloNet analyses^31,41^ are powerful tools to detect ancestral gene flow, such as the prominent signal between the Asiatic black bear and the ancestor to the American black, brown and polar bears (Fig. 4, Table 1). In fact, gene flow during early ursine radiation from extinct bear species, such as the Etruscan bear or the cave bear is to be expected to leave signatures in genomes of their descendants and thus causing conflict in a bifurcating model of evolution.

**Table 1.** Gene flow detected by the *D_FOIL_* analyses that is based on a five taxon analysis. Note – The table shows the percentage of 100 kb fragments that have a signal of gene flow, and in brackets the absolute number is shown. The rows show these values for different combinations of four bear species with the spectacled bear as an outgroup. The last three columns summarize amount of gene flow. The arrows in the table (=>) indicate the direction of the gene flow, between the respective species for each of the combinations analyzed. For example: between Asiatic black and American black bear the *D_FOIL_* finds 15 – 34 GF that support gene flow (first row). There is much more gene flow in the other direction (second row). Abbreviations: SuB (Sun bear), SlB (Sloth bear), AsB (Asiatic black bear), AmB (American black bear), BrB (Brown bear, Finland), and PoB (Polar bear, Svalbard).

|  | AmB,<br>BrB,<br>SuB,<br>AsB | AmB,<br>BrB,<br>SIB,<br>AsB | AmB,<br>PoB,<br>SuB,<br>AsB | AmB,<br>PoB,<br>SIB,<br>AsB | Average %<br>(As/AmB,<br>SuB/SIB,<br>PoB/BrB) | Average %<br>(BrB, AmB,<br>SuB/SIB,<br>AsB) | Average %<br>(PoB, AmB,<br>SuB/SIB,<br>AsB) |
| --- | --- | --- | --- | --- | --- | --- | --- |
| **AsB => AmB** | 0.07% (15) | 0.08% (18) | 0.14% (31) | 0.15% (34) | 0.11% |  |  |
| **AmB => AsB** | 1.22% (270) | 1.42%(313) | 2.06% (454) | 2.48% (547) | 1.80% |  |  |
| **Su/SIB => AmB** | 0.02% (4) | 0.01% (1) | 0.02% (5) | 0.02% (5) |  | 0.01% | 0.02% |
| **AmB => SuB/SIB** | 0.02% (4) | 0.01%(1) | 0.02% (4) | 0.01% (2) |  | 0.01% | 0.01% |
| **SuB/SIB => BrB/PoB** | 0.07% (16) | 0.02%(4) | 0.02% (5) | 0.02% (5) |  | 0.05% | 0.02% |
| **BrB/PoB => SuB/SIB** | 0.03% (6) | 0.01%(1) | 0.02% (5) | 0.01% (3) |  | 0.02% | 0.02% |
| **AsB => BrB/PoB** | 1.25% (276) | 1.02%(225) | 0.46% (101) | 0.29 (64) |  | 1.14% | 0.37% |
| **Br/Po => As** | 9.70% (2159) | 11.0% (2415) | 8.37% (1846) | 9.02 (1989) |  | 10.4% | 8.69% |
| **BrB/PoB, AmB <=> SuB/SIB** | 0.10% (23) | 0.06 (14) | 0.20% (44) | 0.11% (25) | 0.12% |  |  |
| **BrB/PoB, AmB <=> AsB** | 32.2% (7098) | 32.0% (7060) | 46.3% (10214) | 45.8% (10108) | 39.1% |  |  |

## Speciation as a selective rather than an isolation process

There is no question that bears are morphologically, geographically and ecologically distinct and they are unequivocally accepted as species even by different species concepts^54^. Yet, our genome-wide analyses identify gene flow among most ursines, making their genome a complex mosaic of evolutionary histories. Increasing evidence for post-speciation gene flow among primates, canines, and equids^22,23,55^ suggests that interspecific gene flow is a common biological phenomenon. The occurrences of gene flow and to a lesser extent ILS, of which a fraction in the phylogenetic signal cannot be excluded, suggest that the expectation of a fully resolved bifurcating tree for most species might be defied by the complex reality of genome evolution. Recent genome-scale analyses of basal divergences of the avian^56^, insect^57^ and metazoan^58^ tree share the same difficulties to resolve certain branches as observed for mammals^59^. Detecting gene flow for these deep divergences is difficult and therefore most of the reticulations and inconsistent trees have been so far attributed to ILS^60^.

The recent discoveries of gene flow by introgressive hybridization in several mammalian species^22,23,55,61^ and in bears over extended periods of their evolutionary history have a profound impact of our understanding of speciation. If, in fact gene flow across several species is frequent, and can last for several hundred-thousand years after divergence, evolutionary histories of genomes will be inherently complex and phylogenetic incongruence will depict this complexity. Therefore, speciation should not only be viewed as achieving genome-wide reproductive isolation but rather as selective processes that maintain species divergence even under gene flow^62^.

## Materials and Methods

### Genome sequencing, mapping and creation of consensus sequences

Prior to sampling and DNA extraction and evolutionary analyses, pedigrees from zoo studbooks and appearance of the individuals confirmed that these individuals are not hybrids ( Supplementary Fig. 3). DNA extraction from opportunistically obtained tissue and blood samples was done in a pre-PCR environment on different occasions to avoid confusion by standard phenol/chloroform protocols and yielded between 1 to 6 μg DNA for each of the six bear individuals (S1 Data 1.1). Paired end libraries (500 bp) were made by Beijing Genome Institute (BGI) using Illumina TrueSeq and sequencing was done on Illumina HiSeq2000 resulting in 100 bp reads. Raw reads were quality-trimmed by Trimmomatic^63^ with a sliding window option, minimum base quality of 20 and minimum read length of 25 bp. The assembled polar bear genome^28^ was used for reference mapping using BWA version 0.7.5a^64^ with the BWA-MEM algorithm on scaffolds larger than 1Mb. Scaffolds shorter than 1 Mb in length were not involved in the mapping and analyses, due to potential assembly artefacts^65^ and for reducing the computational time in downstream analyses. Duplicate Illumina reads were marked by Picard tools version 1.106 (http://picard.sourceforge.net/) and the genome coverage was estimated from Samtools version: 0.1.18^66^.

Freebayes version 0.9.14-17^67^ called Single Nucleotide Variants (SNVs) using the option of reporting the monomorphic sites with additional parameters as -min-mapping-quality 20, -minalternate-count 4, -min-alternate-fraction 0.3 and -min-coverage 4. A custom Perl script created consensus sequences for each of the mapped bear individuals from the Variant Call Format (VCF) files, keeping the heterozygous sites and removing indels. In order to complete the taxon sampling of the ursine bears, reads from six previously published genomes (S1 Data 1.1) were downloaded on the basis of geographic distribution, availability and sequence depth and SNVs were called as described above. For the two high coverage (>30X) genomes, SNVs calling parameters were set as one-half of the average read depth after marking duplicates. Genome error rates^68^ were calculated on the largest scaffold (67 Mb) for all bear genomes, confirming a high quality of the consensus sequences. For additional details see Supplementary Methods and Supplementary Fig. 22.

## Data filtration, simulation of sequence length and topology testing

The next step was to create multi-species alignments that could be used for further phylogenetic analysis from all 13 bear individuals included in this study. In order to create a data set with reduced assembly and mapping artefacts, genome data was masked for TEs and simple repeats^22^ using the RepeatMasker^69^ output file of the reference genome (polar bear) available from http://gigadb.org/^28^. Since the polar bear reference genome RepeatMasker output file did not contain the simple repeat annotation, we repeatmasked the polar bear reference genome with the option (-int) to mask simple repeats. Then all the bear genomes were masked with the help of bedtools version 2.17.0^70^ and custom Perl scripts. Non-overlapping, sliding window fragments of 100 kb were extracted using custom perl scripts together with the program splitter from the Emboss package^71^ from scaffolds larger than 1 Mb (Supplementary Fig. 1), creating a dataset of 22,269 GFs from 13 bear individuals. In addition, heterozygous sites, and repeat elements and ambiguous sites were all marked “N “ and removed using custom Perl scripts. An evaluation of the minimum sequence length of GFs needed for phylogenetic analysis was done by estimating how much sequence data is needed to reject a phylogenetic tree topology with using the approximate unbiased, AU test^72^. This is important, because only sufficiently long sequences can differentiate between alternative trees with statistical significance. The evaluation was done in two separate analyses: (a) with a simulated data set and (b) on a data set of 500 random GFs. For additional details see Supplementary Methods.

## Phylogenetic tree analysis using Genomic Fragment (GF) data

For phylogenetic analysis, all GFs with length <25 kb were removed from the initial 22,269 GFs resulting in a data set consisting of 18,621 GFs (mean sequence length of 46,685 bp and standard deviation of 9,490 bp). The dataset was then used to create a coalescent phylogenetic species tree. First the selected GFs were used to create individual maximum likelihood (ML) trees using RAxML version 8.2.4^73^. The best fitting substitution model was selected on 10 Mb of genomic data using jModelTest 2.1.1^74^ available in RAxML version 8.2.4^73^ and applied to all ML analyses. From 18,621 ML trees, Astral program^30^ constructed a coalescent species tree. For bootstrap support of the coalescent species tree, GF were bootstrapped 100 times, generating a total of 1,862,100 ML trees. The bootstrapped ML-trees and the coalescent species tree were used as input in Astral^30^ using default parameters to generate bootstrap support. The consense program in Phylip version 3.69^75^ built from 18,621 ML trees a majority rule consensus tree. SplitsTree version 4^76^ created a consensus network from the 18,621 GF ML-trees with various threshold settings (5%, 7%, 10% and 30%), to explore the phylogenetic conflict among the bear species.

## Phylogenetic analysis of nuclear protein-coding genes and mitochondrial genomes

The annotation of the polar bear genome^28^ was used to extract the protein coding sequences (CDS) from the genome that could be used for phylogenetic analysis. The species alignments were complemented by giant panda (*Ailuropoda melanoleuca*, ailMel1) sequences from Ensembl (http://www.ensembl.org/) to provide an outgroup to the analyses. In order to determine orthologous CDS between the polar bear reference genome and the giant panda Proteinortho version 5.06^77^ was used. The CDS of the new bear genomes that corresponded to the polar bear CDS were aligned with MAFFT version 7.154b^78^. Gaps were removed using Gblocks version 0.91b^79^ and a custom perl script removed ambiguous sites. CDS <300 bp were not used for phylogenetic analyses. The best evolutionary model, GTR+G+I^80^, was estimated using jModelTest 2.1.1^74^. A coalescent species tree was constructed with bootstrap support with Astral^30^ from individual CDS using the GTR+G+I model of sequence evolution. In addition, a concatenated analysis of the coding sequence was also done to estimate the concatenated CDS tree. The CDSs were concatenated and the substitution model GTR+G+I was used to create an ML tree with RaxML^73^. A AU topology test was made on the CDS topology using the CONSEL version 1.20^81^ and the species tree (Fig. 2A).

In order to extract the complete mt genomes from Illumina sequence data, the reads for different bear species were mapped to their respective published complete mt genome sequences using BWA version 0.7.5a^64^. Consensus sequences were created using Samtools version: 0.1.18^66^, aligned by MAFFT version 7.154b^78^ to 32 published sequences (Accession numbers see Supplementary Fig. 12), and MrBayes version 3.2.2^82^ was used to create the Bayesian phylogenetic tree using the best fitting GTR+G+I model of sequence evolution. The analysis was run for 4,000,000 generations with a sample frequency of 4,000 with default priors and an arbitrary burn in of 25% of the samples. Convergence was assessed using the average standard deviation of split frequency which reached < 0.01 and potential scale reduction factor close to 1.00.

## Gene flow analysis using *D*-statistics and the *D_FOIL_*-method

The program ANGSD^83^ was used for admixture analysis (*D*-statistics) among the ursine bears using the spectacled bear-Chappari as outgroup. The reads of other bears were mapped to the consensus sequence of the spectacled bear as previously described. In addition, insertion/deletion (indel) realignment was done using GATK version 3.1-1^84^. All possible four-taxon topologies of the bear species including sun bear-Anabell, brown bear-Finland, Brown bear-ABC, Polar bear-2, American black bear, Asiatic black bear, Sloth bear were involved for gene flow analysis using *D*-statistics. A block jackknife procedure (with 10 Mb blocks) with parameters: -minQ 30 and -minMapQ30, was used to assess the significance of the deviation from zero. We also mapped the sun bear-Anabell, the Asiatic black bear and the sloth bear against the giant panda genome (ailMel1) http://hgdownload.soe.ucsc.edu/goldenPath/ailMel1/bigZips/ and repeated the analyses described above on to investigate, if the choice of the outgroup affected our conclusions. In addition, we analyzed the data using *D_FOIL_*-statistics^31^, to detect the signatures of introgression. For this analysis we assumed the coalescent species tree (Fig. 2A) and selected a window size of 100 kb with --mode dfoil as suggested by the authors^31^. Other parameters were left at default.

## Hybridization inference using PhyloNet

A data set of 4,000 random (every fourth) GFs, that are putatively in linkage equilibrium, was created to calculate rooted ML trees with RAxML as described earlier. The trees were pruned to contain one individual of each ursine species plus the ABC- brown bear to reduce computational complexity of the ML analyses. Maximum likelihood networks in a coalescent framework, thus incorporating ILS and gene flow, were inferred using PhyloNet^41,42^ allowing 0, 1 and 2 reticulations in 50 runs and returning the five best networks.

## Estimation of heterozygosity, past effective population size and divergence times

In order to calculate the amount of heterozygous sites as well as their distribution in all the bear genomes, their genomes were fragmented into 10 Mb regions using custom Perl scripts. The number of heterozygous sites was counted using a custom Perl script and plotted as distributions using R. The pairwise sequentially Markovian coalescent (PSMC)^45^ analysis was done to assess past changes in effective population size over time. We used default parameters and 100 bootstrap replicates assuming a generation time for brown and polar bears of ten years, and six years for the other bear species for the PSMC analysis. We selected a mutation rate of 1×10^-8^ changes/site/generation for all species. These parameters were used in previous brown and polar bear^26^ and enable comparability between the studies. A generation time of six years has been shown for the American black bear^85^ and was deemed realistic for the other relatively small-bodied bears. The mutation rate is close to a pedigree-based mutation rate of 1.1×10^-8^ changes/site/generation in humans^86^ that is considered to be typical for mammals.

A well-documented fossil from the giant panda lineage at 12 million years ago (Ma)^20^ with a maximum calibration point of 20 Ma based on mitochondrial estimates^87^ was used to provide the calibration point needed in PAML MCMCtree^88^ to estimate divergence times on 5,151,660 bp coding sequence data. In addition, the divergence time of Tremarctinae was set to 7-13 Ma^89^. Divergence time estimates of Ursinae was based on the occurrence of *U. minimus* at 4.3-6 Ma^90^, and a polar/brown bear divergence was given a range of 0.48-1.1 Ma^8,10,28^. The calibration points were used to estimate divergence times in MCMC tree in PAML^88^ with a sample size of 2,000,000, burn-in of 200,000, and tree sampling every second iteration. Convergence was checked by repeating the analysis again.

## Acknowledgements

We are grateful to Luay Nakhleh (Rice University) for expert help with Phylo-Net analyses, Yichen Zheng for valuable comments on the manuscript and to Jon Baldur Hlidberg (www.fauna.is), and Aidin Niamir for artwork. Blood samples were kindly provided by Ludwig Carsten (Allwetter Zoo Münster), Tim Schikora (Zoo Schwerin), Christian Wenker (Basel Zoo) and Eva Martinez Nevado (Zoo Madrid). This study was supported by Hesse’s funding program LOEWE (Landes-Offensive zur Entwicklung Wissenschaftlich-ökonomischer Exzellenz) and the Leibniz Society.

### Author contributions

A.J. designed the research and obtained funding. A.J. and T.B. collected the data; V.K. and F.L. conducted the analyses; L.K. provided pedigrees and located samples; A.J., V.K., M.P., M.N., F.L., and T.B. interpreted the results; A.J. and V.K. wrote the paper with the help of all authors.

### Completing Financial Interests

The authors declare no competing interests.

### Data submission

The raw reads of the genome sequences have been deposited in the European Nucleotide Archive under the BioProject accession code PRJEB9724.

